# Single-Cell Phenotypic Heterogeneity Shapes Quorum Signaling Dynamics in *Pseudomonas aeruginosa*

**DOI:** 10.1101/2025.05.29.656822

**Authors:** Danielle Gabi-Lange, Vadim Litvinov, Daniel Dar

## Abstract

Quorum sensing (QS) orchestrates collective responses in bacteria. Yet, how individuality arises and functions within a system designed to unify behavior remains poorly understood. Here, we use imaging-transcriptomics to comprehensively profile the QS transcriptional response of the opportunistic pathogen *Pseudomonas aeruginosa* at single-cell resolution. We observe heterogeneity across all stages of QS, from signal production to public goods expression. While most cells cooperate, the extent of public goods contribution varies substantially, primarily due to transcriptional noise. Additionally, cells expressing high levels of a given exoproduct were often enriched for multiple other public goods. Expression of QS signal synthases, particularly in the Las and PQS systems, exhibits extreme cell-cell variability indicative of regulation, and exceeding levels observed in known differentiation into hypervirulent or motile subpopulations. These patterns are dynamic, peaking during early signaling, suggesting that a subset of highly committed cells may influence key decision points in the QS response. We find that cellular memory carried from prior growth cycles shapes QS dynamics via signaling-primed cells but does not influence *de novo* formation of hyper-signaling cells. Differentiation into hyper-signaling subpopulations is also robust to exogenous autoinducers across a wide concentration range, suggesting it is likely mediated by an internal mechanism. Despite variation in QS network outputs between strains, hyper-signaling subpopulations consistently emerge across diverse isolates, pointing to an evolutionarily conserved strategy. Our findings recast QS as a system that embeds a deliberately heterogeneous entry point within an otherwise synchronizing program, offering new insight into how bacterial populations balance cooperation and conflict.

## Introduction

Bacteria use quorum sensing (QS) to orchestrate collective behaviors such as virulence factor production and biofilm formation in a cell density-dependent manner^1,2^. In Gram-negative species, this process typically involves two types of genes: a signal synthase and its cognate receptor^3^. The synthase produces small diffusible signaling molecules, known as autoinducers, which are secreted into the environment and serve as proxies for cell density. This enables cells to collectively regulate production of shared resources (i.e., public goods), such as exoproteases and iron-scavenging siderophores, and effectively coordinate tasks that exceed the capabilities of an individual cell. In traditional models, once an autoinducer threshold concentration is reached, receptor proteins bind their respective autoinducers and act as transcription factors to trigger a sharp QS response across the population. However, recent studies suggest the QS response can follow a graded, rather than step-function, activation, and that cells may participate in a non-uniform manner^4,5^. Although QS has been extensively characterized in diverse species and contexts, a fundamental question remains: how does individuality arise and function within a system designed to synchronize behavior?

Clonal bacterial populations frequently display phenotypic heterogeneity at the single-cell level, which can have major implications for traits such as antibiotic tolerance and virulence^6^. Much of this variability stems from stochastic gene expression and environmental fluctuations—collectively referred to as gene expression ‘noise’^7,8^. However, heterogeneity is not purely passive: bacteria can actively regulate gene expression to restrict specific functions such as motility, adhesion, or virulence to distinct subpopulations^6,9^. This functional diversification supports multicellular-like behavior in otherwise unicellular organisms. By contrast, QS is traditionally viewed as a unifying mechanism that coordinates cells into collective action. Yet, QS imposes metabolic costs on individual cells, producing expensive public goods^10^ and signaling molecules^11,12^. This tension between individual cost and group benefits echoes broader principles in microbial social evolution, where cooperative traits are susceptible to exploitation by non-contributing individuals (“cheaters”)^10^. Importantly, it raises key questions about how coordination is regulated at the single-cell level. Understanding this balance is essential for uncovering how bacterial populations manage cooperation, conflict, and possibly division of labor.

Recent single-cell studies using fluorescent reporters have begun to uncover phenotypic heterogeneity in QS-regulated traits, indicating that individual cells within clonal populations may contribute unequally to shared tasks^13,14^. However, these studies have primarily relied on single-gene measurements, which limits their ability to quantitatively distinguish stochastic expression noise from regulated differentiation. Additionally, they provide limited insight into how the expression of multiple interconnected QS genes is partitioned across individual cells. Thus, the broader architecture and dynamics of QS phenotypic heterogeneity are largely unexplored. Moreover, the mechanisms underlying this variation, as well as its functional consequences and evolutionary conservation, remain poorly understood.

Here, we address these knowledge gaps using advanced single-cell transcriptomics and quantitative analysis of cell-cell variability. Focusing on the opportunistic human pathogen and long-standing QS model organism, *Pseudomonas aeruginosa*, we reveal new insights into how phenotypic heterogeneity shapes cooperative behavior in bacterial populations.

## Results

### Targeted single-cell transcriptomics of P. aeruginosa quorum sensing

*P. aeruginosa* encodes three interlinked QS systems that are hierarchically organized. The Las and Rhl systems use acyl-homoserine lactone (AHL) autoinducers, while the PQS system relies on quinolone-based signals^15^. At the top of the regulatory hierarchy, the Las system—comprising the synthase *lasI* and the receptor *lasR*—initiates the QS cascade, followed by activation of the PQS and Rhl systems. However, a complex network of interactions among the three systems shapes the expression of QS-regulated exoproducts^16^. To resolve the dynamics of this network, which involves dozens of genes governing signal production, sensing, regulation, and downstream responses, we applied parallel and sequential Fluorescence *In Situ* Hybridization (par-seqFISH)^17^. This approach enables multiplexed imaging of bacterial gene expression across dozens of conditions at single-cell resolution. We designed a probe library targeting all known QS synthase and receptor genes, key regulators, and both direct and indirect QS-regulated exoproducts from the Las, Rhl, and PQS systems (Table S1)^18,19^. In addition, we included marker genes reporting on growth, metabolism, stress, and virulence. In total, our panel covered 144 genes, capturing the QS network and broader physiological context within individual cells.

To track density-dependent transcriptional changes, we grew *P. aeruginosa* PAO1 planktonically in rich medium (LB) and collected cells at 11 optical densities (OD₆₀₀), spanning exponential to stationary phase (Fig. 1A). Cells from each timepoint were labeled with a unique 16S rRNA barcode probe (i.e., Ribo-Tag), allowing samples to be mixed and jointly hybridized with a pool of mRNA binding probes (Methods). Fluorescent barcodes and transcript signals were acquired sequentially by combined microfluidics and automated microscopy (Fig. 1A). We demultiplexed 63,972 cells, with an average of 5,815 cells per condition and an estimated misclassification rate of 0.03% (1 in 3,300). To validate that our dataset captures expected QS behavior, we computed the mean expression of each gene at each density (Table S2). Expression levels were highly reproducible across biological replicates (r = 0.99, *p* < 10⁻^10^; Fig. S1). Physiological markers varied in line with known density-dependent shifts, including reduced ribosomal subunit expression and induction of anaerobic metabolism at higher densities (Fig. 1B; Table S2). QS activation followed the expected density-dependent pattern: the *lasI* synthase was induced during mid-exponential phase, followed by the secondary synthases (*rhlI* and *pqsABCDE*) as cells transitioned to stationary phase (Fig. 1C, Table S2). The *lasR* receptor was expressed even at the earliest time points and maintained high expression across nearly all physiological states examined (Fig. 1D, Table S2). The expression of QS-regulated exoproducts (e.g., *cbpD*, *lasB*, *rhlB*) occurred at higher cell densities, consistent with the established regulatory hierarchy (Fig. 1E, Table S2). These results confirm that our single-cell imaging-transcriptomics approach captures both the average QS transcriptional dynamics and concurrent physiological transitions during population growth.

**Figure 1.**
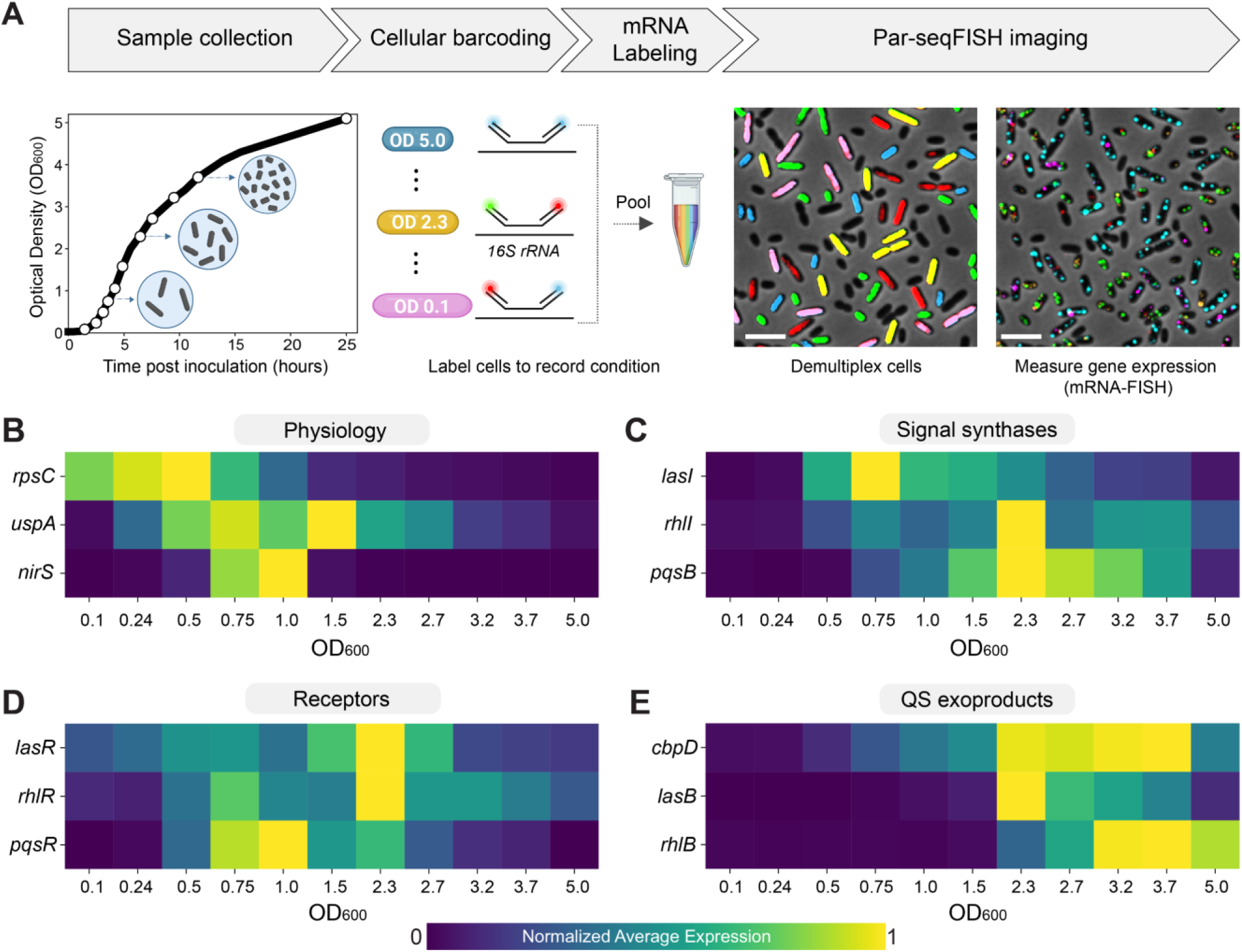
Par-seqFISH imaging of the *P. aeruginosa* QS response. **(A)** Experimental outline, including sample collection, multiplexing, and image acquisition. Cells were collected and fixed across time points as indicated in the growth curve. Samples were mixed and analyzed together using multiplexed par-seqFISH imaging. Representative 16S rRNA barcodes and mRNA FISH signals are shown over phase contrast images. Scale bar = 2 μm. Average expression across each density for genes representing key physiological markers **(B)**, QS synthases **(C)**, receptors **(D)**, and products **(E)**. Expression of each gene is normalized to its maximum across all densities.

### Cooperation is widespread but strongly influenced by high-contributing minority subpopulations

The complexity of the *P. aeruginosa* QS network poses significant challenges for analysis using single-gene reporter assays. Our single-cell transcriptomic data offers an alternative approach, enabling quantification of individual contributions to public goods production across the entire network. We measured the proportion of cells expressing each gene per optical density (Table S2), finding strong agreement for this metric between biological replicates (r = 0.98, *p* < 10^⁻10^; Fig. S2A). At high densities, up to 90% of cells expressed at least one QS-regulated exoproduct gene (e.g., *aprA*, *lasA, cbpD, etc.;* Table S2; Fig. S2B). However, while our mRNA measurements capture active cooperation, they are also sensitive to the global reduction in transcription that occurs as growth slows^20^. This physiological response, together with the short mRNA half-lives of bacteria^21^, constrains the potential number of cells actively expressing any gene at higher densities. Indeed, although some genes are expressed in nearly all cells during rapid growth (e.g., ribosomal protein *rpsC*), no gene was detected in more than 39% of cells by early stationary phase (Fig. S2C; Table S2). By normalizing for this global reduction, we decoupled the effects of cell density and growth-phase physiology (Method) and found near-saturated expression-potential even for individual exoproduct genes (Fig. 2A), consistent with previous single-gene fluorescence reporter studies^4,5,22^.

**Figure 2.**
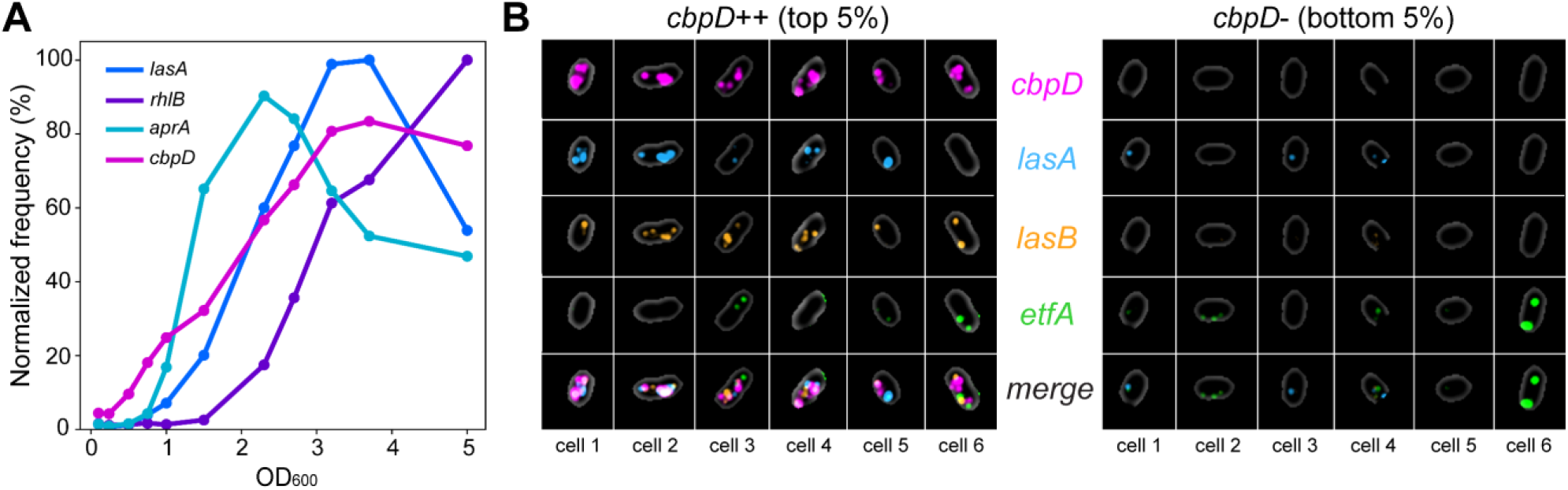
Evaluating the contributions of individual bacteria to the QS response. **(A)** Expression frequency for directly regulated QS exoproducts (*lasA, rhlB, aprA, cbpD*) per optical density. Color codes per gene are shown within the figure. The y-axis shows the expression frequency normalized by the maximum observed frequency per given density. For instance, the maximal frequency observed at OD_600_ of 2.7 was 72% and defined as the normalized 100% (Methods; Table S2). **(B)** Six randomly selected *cbpD*++ (top 5% of *cbpD* expressers) and *cbpD*-cells (bottom 5% of *cbpD* expressers) from an OD_600_ of 3.7. Cells are shown via phase contrast and overlaid with smFISH data for key exoproducts (*cbpD, lasA, and lasB*), as well as a non-QS control gene (*etfA*) that is expressed to similar levels in both groups.

The above frequency analysis indicates that most cells contribute to public goods production; however, it remains unclear whether contribution magnitudes are uniformly or variably distributed across the population. To explore this, we ranked cells based on the expression level of individual exoproduct genes (e.g., *cbpD*) and compared the top and bottom 5% expressing subpopulations (e.g., *cbpD++* vs. *cbpD-*; Methods). Notably, cells expressing high levels of a given exoproduct were often enriched for multiple other public goods. For example, the *cbpD++* subpopulation exhibited up to 5.6-fold higher expression of exoproducts such as *lasA*, *lasB*, and *rhlB* in early stationary phase, along with a 3.6-fold enrichment of PQS synthase genes (Fig. 2B, Fig. S2D; Table S3). These findings suggest that certain subpopulations contribute disproportionately—well beyond their “expected share”—to overall public good abundance. Indeed, in population densities ranging from OD₆₀₀ of 2.3 to 3.7, the *cbpD++* subpopulation accounted for 15.9–18.7% of the total expression of key exoproduct genes, more than threefold higher than expected based on their frequency. Despite their elevated investments, we found no evidence that these highly engaged cells incur immediate fitness costs, as key growth marker genes (e.g., *rpsC*) were expressed at levels comparable to the *cbpD*-subpopulation. We extended the same “top 5 % vs. bottom 5 %” analysis to higher-order regulators and find that *lasR*++ cells show elevated *lasI, rhlI, rhlR* up to 6-fold, while *rhlR*++ cells boost *rhlI* and *rhlB* by up to 4-fold (Table S3). These data show that receptor mRNA levels can report on receptor activity. Thus, co-existing subpopulations are differentially engaged in QS actions.

### Systematic identification of hypervariably expressed genes

Our data suggests that individual subpopulations can contribute disproportionately to QS. While it is tempting to speculate that bacteria actively regulate this heterogeneity, an alternative explanation is that it may arise passively from the baseline stochasticity inherent in bacterial gene expression^7^. To distinguish cases in which phenotypic heterogeneity significantly exceeds such background noise— and may therefore reflect regulatory control—we systematically analyzed cell-cell variability across all genes and conditions. We employed two complementary measures: the coefficient of variation (CV)^23,24^ and a new metric we developed called the Tail Expression Ratio (TER). TER quantifies the breadth of the expression distribution by comparing its extremes (tails), defined as the ratio of mean expression in the top and bottom 5% of cells with non-zero expression (Methods).

Cell-cell expression variability is highly affected by gene mean expression levels^23–25^. We therefore plotted both CV and TER against mean expression for each gene-condition pair, fitting regression lines to model the baseline expression noise (Methods). Deviations from these trends were calculated to obtain the CV and TER residual variability (CVR and TERR, respectively; Table S4), which were integrated into a single Expression Variability Index (EVI; Methods). Importantly, EVI scores showed no correlation with mean expression (r = –0.001; Fig. S3A), confirming effective decoupling of variability from expression level. Moreover, these scores were highly reproducible across biological replicates (r = 0.98; *p* < 10⁻^10^; Fig. S3B).

To identify hypervariable genes, we ranked gene-condition pairs by their EVI scores and selected the top 5% data points (Methods). Notably, *fliC*, encoding the flagellin protein, consistently emerged among the most hypervariable genes in our dataset (Fig. S3C-E; Table S4), in agreement with previously described differentiation into motile and non-motile subpopulations^17,26^. Similarly, our approach also highlighted *exsA*, the master regulator of the type III secretion system (T3SS), which drives the formation of hypervirulent subpopulations^27^ (Table S4). Also among the most hypervariable genes were *algU* and *mucA*, encoding co-transcribed sigma and anti-sigma factors, respectively, that regulate alginate biosynthesis^28^ (Table S4). These examples outline the power of this quantitative approach and provide a foundation for assessing the source of variability in QS-related genes.

### Conserved differentiation into specialized QS signaling subpopulations

Building on the above variability analysis, we next focused on QS components to explore how cell-cell expression variability shapes this key communication system. Strikingly, among all expressed gene-condition pairs in our experiment (n = 782), the *lasI* and *pqsB* genes—encoding synthases of the Las and PQS systems, respectively—exhibited the highest expression variability scores (>99.6th percentile; Fig. 3A–F; Table S4). Additional PQS synthase genes were also detected as hypervariable (Table S4). Furthermore, the *lasI* and *pqsB* EVI score peaks coincide precisely with optical densities corresponding to QS system activation, followed by a marked reduction in heterogeneity as cell density increases (Fig. 3G). This dynamic pattern supports a model in which QS signal production is initially driven by a subset of highly committed cells, with subsequent recruitment of the broader population as signal dissemination progresses. In contrast, *rhlI*, the synthase of the Rhl QS system, displayed expression variability indistinguishable from background noise (Figs. 3G and S3F–H). This divergence between synthase genes highlights distinct regulatory architectures across QS subsystems.

**Figure 3.**
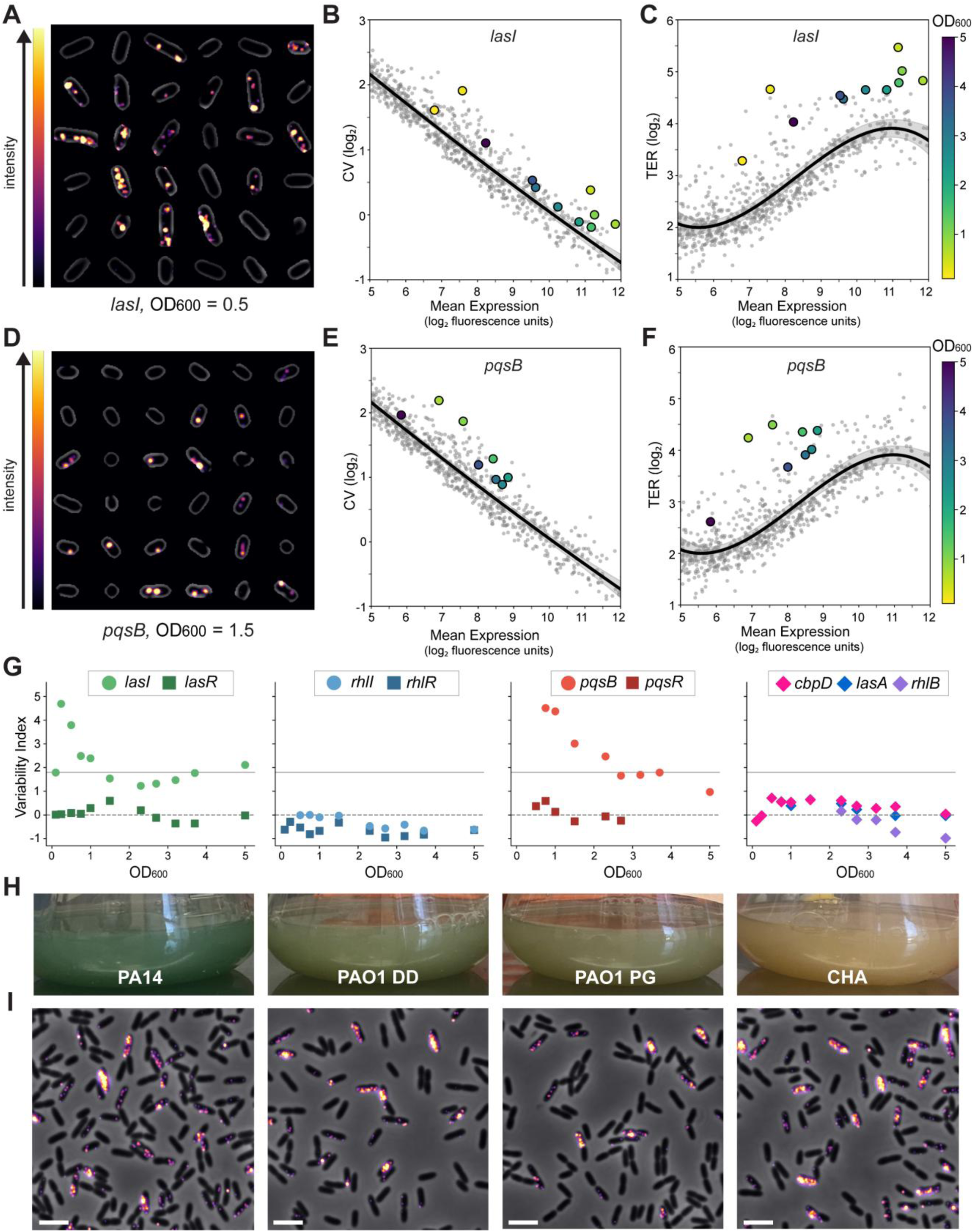
Excess variability analysis reveals specialized signaling subpopulations. **(A)** Phase contrast images of randomly selected cells from an OD_600_ of 0.5 overlaid with smFISH data for *lasI*, with smFISH signal intensity represented using the “inferno” colormap. **(B–C)** Log_2_ transformed CV and TER values plotted against Log_2_ mean expression (arbitrary fluorescence units). Gray points represent expression measurements for all genes across all densities. Black lines indicate regression fits with gray shading for 95% confidence intervals (Methods). The *lasI* data points are highlighted and colored by density as indicated. **(D-F)** Show the same analysis as in panels *(A–C)*, but highlighting *pqsB*, with images in *(D)* showing randomly selected cells from an OD_600_ of 1.5. **(G)** The Expression Variability Index (EVI) plotted against density (OD_600_) for synthase (circles) and receptor (squares) genes associated with the Las (green), Rhl (blue), and PQS (red) QS systems, as well as QS-regulated exoproducts (*cbpD, lasA, rhlB;* diamonds). Dashed and solid horizontal lines indicate EVI values of 0 (no excess heterogeneity) and the cutoff for the top 5% of EVI values in the dataset (1.82), respectively. **(H)** Varying levels of pigmented products (e.g., pyocyanin) after 24 hours of growth for four different *P. aeruginosa*. **(I)** Phase contrast images of cells at OD_600_ of 0.24 overlaid with *lasI* smFISH signal (from left to right; PA14, PAO1 DD, PAO1 PG, and CHA). Scale bar = 2 μm.

Notably, while the Las and PQS signal synthases exhibited variability exceeding even that of established heterogeneous genes (e.g., *fliC* and *exsA*), expression of their cognate receptors—as well as the majority of QS-controlled exoproducts—remained within baseline noise levels (Fig. 3G; Figs. S3I–K; Table S4). To characterize the transcriptional signatures of these hyper-signaling subpopulations, we compared the top 5% vs. bottom 5% of *lasI*-expressing cells (*lasI++*). This analysis revealed significant enrichment for *rsaL*, a direct negative regulator of *lasI* transcription, which is expressed from a shared bidirectional promoter^29^ (Table S3). This *rsaL* upregulation was density-dependent, peaking with a 24-fold increase at OD₆₀₀ = 0.5 and gradually declining to just 2-fold by OD₆₀₀ = 3.2, indicating a decoupling of the RsaL-mediated negative feedback with activation of the QS response. These patterns suggest that variability in QS signaling is tightly regulated, while heterogeneity in public goods production likely arises through passive, stochastic processes.

To assess the evolutionary conservation of this phenomenon, we compared four diverse *P. aeruginosa* isolates: two PAO1 lab strains acquired from different sources, the hypervirulent UCBPP-PA14 strain, and the clinical CHA isolate (Methods). Of note, these strains exhibited significant differences in pigment accumulation (e.g., pyocyanin) under identical growth conditions, indicative of divergence in their QS-regulated outputs as previously described^30^ (Fig. 3H). However, despite these differences, *lasI* single-cell expression patterns were remarkably conserved across isolates (Fig. 3I). Taken together, these findings reveal the presence of conserved subpopulations specialized in QS signaling, suggesting an evolved and potentially regulated differentiation that may underpin cooperative behaviors in *P. aeruginosa*.

### Hypervariability in QS signaling is independent of cellular memory and autoinducer levels

The above analyses uncovered subpopulations specializing in either Las or PQS signaling. However, the mechanisms driving their differentiation remain unclear. In our experiments, cultures were grown overnight, then washed and diluted 1:100 into fresh media (hereafter termed the “D100 experiment”). During this culturing step, cells reached stationary phase, where QS was fully activated, raising the possibility that subpopulations “primed” for QS activation or signaling may persist into the next growth cycle. Conceptually, such cellular memory could act through two non-exclusive mechanisms: in a lineage-based model the same cells (or their descendants) activated in the previous cycle maintain a hyper-signaling state. Alternatively, in an environmental-priming model, activated cells from the prior cycle shape the extracellular context (e.g., through signal production), thereby biasing new cells toward hyper-signaling. Understanding the potential role of either such form of “memory” is essential for elucidating the origins of phenotypic heterogeneity in QS and potentially in other contexts^31^.

We therefore performed another experiment in which cultures were inoculated at an ultralow dilution (1:10,000; hereafter termed the “D10000 experiment”). In this setup, cells underwent approximately seven additional doubling events before reaching the same starting density as in the D100 experiment. These successive divisions in low-density conditions are expected to dilute and degrade residual proteins, thereby erasing the potential effects of stationary-phase memory (Fig. 4A). We collected D10000 samples at the same optical densities as in our original D100 experiment and profiled 58,528 cells across this matched density range using par-seqFISH (Fig. S4A; Methods). As in the D100 experiment, we observed strong agreement in mean expression, expression frequency, and EVI scores between D10000 biological replicates (Fig. S4B-D).

**Figure 4.**
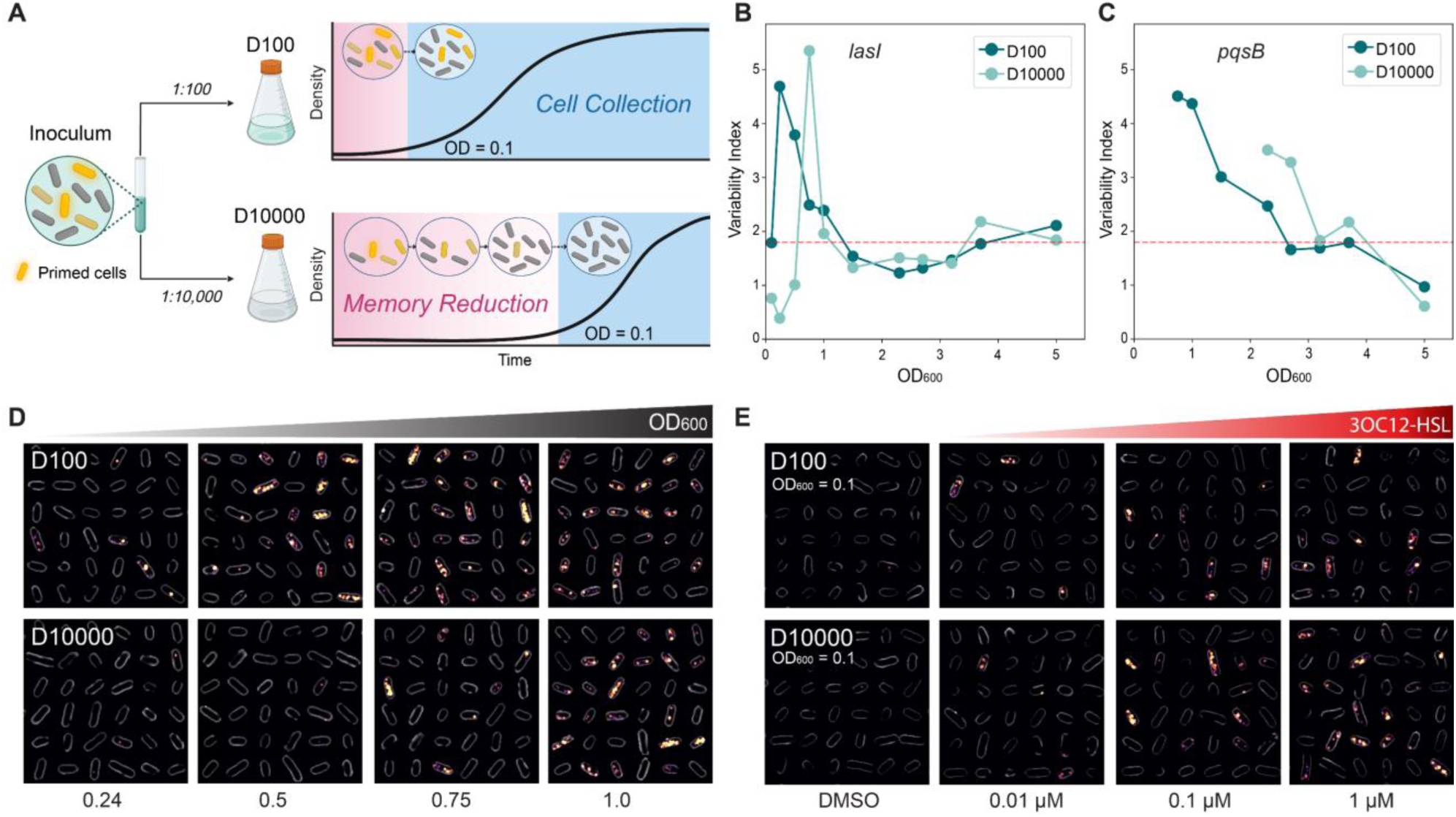
Signaling subpopulations are independent of cellular memory and signal concentration. **(A)** Schematic illustration of the memory erasure experiment. Cultures carrying cells with differential stationary-phase primed cells (yellow cells) are diluted either 1:100 (D100) or 1:10,000 (D10000) into fresh media. Memory is reduced by division-based dilution and protein degradation in D10000 experiment until the culture reaches the collection stage (blue background; OD_600_ = 0.1). **(B-C)** Variability index shown for the *lasI* and *pqsB* signaling genes across density in D100 (dark turquoise) and D10000 (light turquoise) samples as indicated in the legends. **(D)** Randomly selected D100 and D10000 cells from low-density timepoints (OD_600_ of 0.24-1.0) shown with phase contrast and overlaid with *lasI* smFISH data. **(E)** Randomly selected D100 and D10000 cells from an OD_600_ of 0.1 treated with 0.01-1µM exogenous 3OC12HSL or DMSO control. Cells are shown with phase contrast and overlaid with *lasI* smFISH data.

Comparing the D100 and D10000 experiments, we found high correlation in both mean expression (r = 0.94; *p* < 10⁻^10^; Fig. S4E) and gene variability metrics (r = 0.85; ; *p* < 10⁻^10^; Fig. S4F), indicating that memory erasure had a minimal impact on most genes. In contrast, both *lasI* and the PQS synthesis genes (e.g., *pqsB*) were highly affected by “memory loss”, showing substantial differences in expression (up to 24-fold for *lasI* at OD_600_ = 0.5, up to 20-fold for *pqsB* at OD_600_ = 1.5; Fig. S4E). However, upon closer examination, we found these differences were due to a temporal delay in induction in the D10000 experiment (Fig. S5). Notably, both *lasI* and *pqsB* eventually reached comparable maximal expression levels (Fig. S5A-B). A similar shift was also detected in Rhl signaling and in QS exoproduct expression, albeit with milder differences (Fig. S5C-D). To evaluate whether these differences were mediated by primed cells enhanced for signaling or sensing, we supplemented D100 and D10000 cultures with increasing concentrations (0.1–1 µM) of the Las signal, N-3-oxo-dodecanoyl homoserine lactone (3OC12-HSL). (Methods). In both D100 and D10000, *lasI* expression increased in a 3OC12-HSL dose-dependent manner and to a similar magnitude, indicating that both cultures were responsive to a comparable degree at these autoinducer concentrations (Fig. S6). Only the DMSO controls reproduced the memory-dependent temporal shifts in *lasI* expression, indicating differences in autoinducer concentration underlie this phenotype (Fig. S6). Thus, these data are consistent with model in which signaling-primed cells acquired at the inoculation stage accelerate autoinducer accumulation and subsequent QS activation, suggesting that single-cell phenotypic memory shapes population-level QS dynamics.

Considering cell-cell variability, we found that both *lasI* and *pqsB* displayed comparable levels of heterogeneity between the D100 and D10000 experiments, differing only in the timing of their induction (Fig. 4B-D, Table S4). Thus, the emergence of Las- and PQS-hyperactive subpopulations appears to occur independently of the initial frequency of primed cells and autoinducer accumulation rates. Indeed, we found that *lasI* expression heterogeneity was largely unaffected by the addition of exogenous autoinducer across a 100-fold concentration range, with clear subpopulation emergence (Fig. 4E). Together, these findings suggest that Las signaling subpopulations—and potentially PQS subpopulations as well—arise independently of both inoculum history and external signal levels, pointing toward an internally regulated mechanism of differentiation.

## Discussion

In this study, we observed heterogeneity across all stages of QS, from signaling to public goods production. However, our data suggest that the sources of heterogeneity differ at each stage: hyper-signaling subpopulations likely form through a regulated process, while differential public good production arises from stochastic events. QS signal synthase genes, particularly Las and PQS, showed remarkable variability—surpassing even *fliC* and *exsA*, which are associated with well-characterized differentiation into motile and hypervirulent subpopulations, respectively, and where heterogeneity has been proposed to confer adaptive benefits^17,26,27^. In contrast, the Rhl synthase gene, as well as receptors of all three QS systems, displayed variability consistent with basal noise, indicating system and network node-specific effects, in agreement with recent reports^22^. These patterns were dynamic, peaking at early stages of signaling, suggesting that subsets of highly committed cells disproportionately influence key decision points. While QS networks and outputs can vary between strains^30^, hyper-signaling subpopulations consistently emerge in four isolates, pointing to an evolutionarily conserved differentiation strategy in *P. aeruginosa*. These findings align with evidence of signal synthesis specialization in other species, underscoring the importance of understanding its function and underlying mechanisms^13,14,32–34^.

Phenotypic heterogeneity is generally attributed to physiological, environmental, and genetically encoded regulation^6^. Here, we focused on cellular memory and autoinducer concentrations, representing physiological and environmental influences, respectively. “Memory” refers to the physiological state inherited from past growth cycles, a concept previously explored in the contexts of lag-phase exit and antibiotic tolerance^35,36^ . We find that memory shapes QS temporal dynamics at the population level, consistent with recent theoretical predictions^31^. Our data suggest that this effect is mediated by a subset of cells primed for signaling, bridging individual heterogeneity and collective behavior. However, neither memory nor external autoinducers significantly influenced the *de novo* emergence of hyper-signaling subpopulations, suggesting a genetically encoded mechanism. Supporting this view, recent work implicates the QS repressor RsaL in *lasI* expression heterogeneity^22^. Our data shows that *rsaL* enrichment in *lasI*++ cells is correlated with *lasI* variability, consistent with this model. Regulation of QS signaling differentiation may be a widespread feature of bacterial communication. For example, in *Sinorhizobium meliloti*, variability in *sinI* expression is shaped by the abundance and affinity of its cognate receptor, SinR^37^. Identifying the promoter features, feedback topologies, and chromosomal contexts that allow Las and PQS—but not Rhl—synthase genes to exhibit such pronounced variability will clarify how bacteria achieve gene-specific control over heterogeneity.

Why do bacteria favor hypervariability in QS signaling? One explanation is that the benefits of signal synthesis are realized under uncertain future conditions, making it an ideal substrate for bet-hedging. Another possibility is that heterogeneity enables division of labor, restricting the metabolic cost of signal production to a subset of cells. This may also protect the population from cheaters that exploit diffusible autoinducers, which function as public goods, while preserving coordinated group behavior^14^. Both concepts hinge on signal synthesis being metabolically costly. Indeed, recent studies show that QS signal production imposes a significant fitness burden in both heterologous systems like *E. coli*^11^ and native *P. aeruginosa*^12^, under nutrient limitation. Consistent with these findings, in the LB-rich conditions used in our experiments we observed no transcriptomic evidence of a growth disadvantage in Las or PQS hyper-signaling subpopulations. Extending these analyses to nutrient-limited and more environmentally relevant media will be necessary to evaluate the metabolic burden at single-cell resolution. While early estimates of signaling costs focused on ATP expenditure^38^, it has since been shown that the cost of Las signaling arises from depletion of S-adenosylmethionine^11^. However, metabolite depletion costs may only manifest after synthase expression, highlighting the importance of complementing transcriptomic analyses with live-cell imaging.

The results of our single-cell analysis suggest two distinct cost-management strategies may operate at different tiers of the QS network. For low-density signaling, an actively regulated bet-hedging strategy seems to dominate. Only a small cohort of *las*/*PQS* hyper-signalers bear any potential early expenses but accelerate auto-inducer build-up to pull the population past the activation threshold. Under this design, should growth stall, the cost is confined to that minority. At higher densities, where quorum is reached, a more passive load-skewing pattern governs QS-regulated exoproducts. In principle, late-stage costs can be buffered in two ways: different subpopulations can specialize in producing specific products, or one minority subpopulation can contribute more of multiple products. While previous studies reported variability in *P. aeruginosa* QS exoproduct expression, it was not possible to differentiate between these possibilities using their single-gene fluorescent reporters^4,5,39^. Our multiplexed imaging resolves this question, supporting the latter option by showing that hyper-cooperating minorities are co-enriched for multiple products rather than specialize. These patterns are generated by intrinsic transcriptional stochasticity, allowing most cells to invest modestly while overall output remains high. This load-skewing contrasts with the clearer division of labor seen in the *Vibrio harveyi* QS regulon (bioluminescence versus protease production)^40^, suggesting bacteria may evolve different strategies. In summary, we hypothesize that *P. aeruginosa* QS pairs a regulated minority-seeding mechanism to limit early commitment risk at the QS signaling gate with a stochastic load-skewing that may alleviate the metabolic cost of exoproduct synthesis at high densities when resources are scarce.

The experiments in this study were conducted in well-mixed cultures, which minimize the influence of microenvironmental heterogeneity and enable precise quantification of cell-cell variability. This allowed us to isolate intrinsic sources of heterogeneity in the QS response, independent of spatial effects. However, spatial structure plays critical roles in more natural settings such as biofilms, surface-associated growth, or infection. The emergence of distinct signaling subpopulations in these contexts may offer additional advantages for communication and diffusion sensing^41,42^. A key question is whether and how a minority of signal-specialist cells can continue to steer group behavior in spatially structured settings. Addressing this will clarify how spatial organization influences communication and cooperation within bacterial populations^43^.

Through this single-cell transcriptomics study, we demonstrate that the *P. aeruginosa* QS network efficiently synchronizes cells for collective behavior yet is structured around deliberately heterogeneous regulatory nodes that are robust to physiological and environmental variation. In recent years, microbial single-cell and spatial transcriptomics have advanced rapidly^44–48^. These approaches will be essential for uncovering how phenotypic diversity is leveraged to manage cooperation and conflict across bacterial species and ecological contexts.

## Methods

### Bacterial strains and growth conditions

The *P. aeruginosa* PAO1 strain was supplied by the Eldar lab^49^ and was used for all main experiments in this study. This strain was originally acquired from Peter Greenberg’s lab and was thus named *P. aeruginosa* PAO1 PG. For strain comparisons, we included the CHA strain^50^ and two reference *P. aeruginosa* isolates obtained from the DSMZ (German Collection of Microorganisms and Cell Cultures): PAO1 (DSM 19880; termed PAO1 DD) and PA14 (DSM 19882). Bacteria were grown aerobically with shaking at 250 rpm in liquid LB Miller medium (tryptone 10.0 g L⁻¹; yeast extract 5.0 g L⁻¹; NaCl 10.0 g L⁻¹) or on LB agar plates at 37°C. For the time series collections, an overnight LB culture was washed twice using fresh LB to remove residual autoinducers and then diluted 1:100 (D100) or 1:10,000 (D10000) into fresh medium. The cultures were grown at 37°C with shaking at 250 rpm and sampled at various time points based on their optical density (OD_600_). Samples were collected in an OD_600_ range of 0.1-5.0 and immediately fixed by direct addition of paraformaldehyde (PFA; Electron Microscopy Science, 15174) to a final concentration of 2%. Fixation was performed on ice for 1.5 hours, followed by three wash steps with 1X PBS. Fixed samples were resuspended in 70% EtOH and incubated at −20°C overnight to permeabilize the cells.

For the exogenous AHL experiment, we supplemented either D100 or D10000 cultures with 0.01, 0.1, or 1 µM 3OC12-HSL (Sigma-Merck, O9139). DMSO only treatment was used as a negative control. D100 cultures were inoculated at an initial OD₆₀₀ of 0.03 into LB medium containing exogenous 3OC12-HSL while D10000 cultures, initially diluted 100-fold lower, were pre-grown to an OD₆₀₀ of 0.03 before exogenous 3OC12-HSL was added. The cultures were incubated at 37°C with shaking at 250 rpm and sampled at optical densities of OD₆₀₀ = 0.1, 0.24, and 0.5. Collected samples were fixed and permeabilized as described above.

### Par-seqFISH probe design and library generation

Probes were designed as previously described^17^. Briefly, primary probes targeting mRNAs were designed as 30-nucleotide (nt) sequences with a GC content between 45% and 65% from the *P. aeruginosa* PAO1 genome (assembly ASM676v1). Probes containing more than four consecutive identical nucleotides were discarded. The remaining candidate probe sequences were aligned to the reference genome using BLAST, and any probe with off-target binding of 18 or more nucleotides was removed. Negative control probes were selected using the P1 phage genome (NC_005856.1) and processed using the exact same criteria against the PAO1 genome. Non-overlapping gene probes were randomly selected and flanked by overhangs containing two repeats of a secondary hybridization sequence, complementary to a designated fluorescent readout probe^17^. A library of 2,988 primary probes targeting 150 *P. aeruginosa* genes and three negative controls was designed. Probes incorporated sequences for PCR amplification, readout binding sites, and a T7 promoter sequence for amplification as previously described^17^, with the only difference of using two rather than four readout binding sites per probe.

The Twist oligonucleotide pool was resuspended to 1 ng µL⁻¹ in 10 mM Tris-HCl (pH 8.0). Probes were amplified by a two-step PCR using KAPA HiFi HotStart ReadyMix: 12 cycles followed by 9 cycles, with a 67.5°C annealing temperature and standard KAPA HiFi denaturation/extension steps (forward primer: 5′-TTTCGTCCGCGAGTGACCAG-3′; reverse primer: 5′-GCATCCCGACATGGACGTTG-3′). 5 µL of the first reaction seeded the second. Four secondary reactions were pooled and purified on a QIAquick column, yielding high-concentration DNA that showed a single band of the expected length on an Agilent DNA TapeStation. 3 µg of purified DNA were transcribed *in vitro* with the HiScribe T7 kit (NEB, E2040S) according to the manufacturer’s short-transcript protocol (8–12 h, 37°C). 5 µL of RNA was column-purified and quantified, and the remaining 95 µL was immediately reverse-transcribed with Maxima H-Minus reverse transcriptase (ThermoScientific, EP0753) using the forward amplification primer (50°C for 2 h; 55°C for 2 h). RNA templates were removed by alkaline hydrolysis followed by ethanol precipitation, and the resulting single-stranded DNA probes were resuspended in 180 µL nuclease-free water and stored at −20°C for hybridization experiments.

### Coverslip functionalization

Coverslips were cleaned with 100% ethanol and dried for 5 minutes at 90°C. To enable acrylamide binding, coverslips were functionalized with Bind-Silane (3-(Trimethoxysilyl)propyl methacrylate; Sigma-Aldrich, M6514-25ML). Coverslips were immersed in 1% Bind-Silane prepared in 10% (v/v) acidic ethanol (pH 3.5) for 1 hour at room temperature. After incubation, coverslips were washed three times with 100% ethanol to remove excess Bind-Silane and dried for 30 minutes at 90°C. To enhance bacterial adherence, one side of each coverslip was coated with 50% poly-D-lysine solution (Sigma-Aldrich, P7280-5MG) and incubated overnight at room temperature in the dark. The following morning, excess poly-D-lysine was thoroughly rinsed off with nuclease-free water. Coverslips were dried with compressed air and were prepared fresh for each experiment.

### Par-seqFISH primary hybridization and multiplexing

Primary hybridizations were performed using 2 × 10⁸ fixed and permeabilized cells per reaction. Cells were pelleted and resuspended in 20 μL of probe solution (probes in nuclease-free water), then mixed with 30 μL of pre-warmed primary hybridization buffer composed of 50% formamide (Thermo Fisher, AM9342), 10% dextran sulfate (Sigma-Aldrich, D8906), and 2X saline-sodium citrate (SSC) (Gibco LifeTech, 15557-036). Reactions were mixed by gentle pipetting and incubated for 16–48 h at 37 °C, as indicated. After hybridization, cells were washed twice with 75 μL of 55% formamide wash buffer containing 55% formamide, 0.1% Triton X-100 (Sigma-Aldrich, 93443), and 2X SSC, followed by a 30-minute incubation in fresh wash buffer at 37 °C to remove nonspecific probe binding. Cells were then washed twice with 100 μL of 2X SSC.

For par-seqFISH multiplexed reactions, independent fixed samples were hybridized individually with 40 nM of unique 16S rRNA labels for 16 hours. Individual reactions were washed well as described above and then pooled into a single mixture. The pool of barcoded cells was hybridized for 48 hours in a primary reaction containing 10 μL of the amplified gene probe library. In addition, the reaction included 40 nM of a reference 16S rRNA probe to define the cellular rRNA signal per cell for multiplexing purposes (described below). A control sample was resuspended in a mock hybridization lacking probes (i.e., no-library control). The final cell mixture was suspended in 30 μL of 2X SSC and spiked with 0.5 μL of the no-library control cells. Then, 5 μL of this cell suspension was spotted at the center of a functionalized coverslip, incubated at room temperature for 5 minutes to allow cells to adhere to the surface, and centrifuged sequentially (500 × *g* for 5 minutes, followed by 3200 × *g* for 5 minutes) to create a uniform, dense monolayer. Cells were subsequently immobilized using a polyacrylamide hydrogel. A 4% (19 : 1) acrylamide/bis-acrylamide (bioRad, 1610154) mix was prepared by combining 66.4 µL 30 × monomer stock, 30 µL 1 M Tris-HCl, 30 µL 5 M NaCl, and 372 µL ultrapure water, vortexing, and degassing the solution on ice for 15 minutes using N_2_. Immediately before casting, polymerization was initiated with 1 µL freshly made APS (50 mg in 200 µL water) and 1 µL N,N,N′,N′ -Tetramethylethylenediamine (TEMED). 10 µL of the reactive mixture were spread over the cell monolayer, covered with a clean slide, and placed inside a sealed container that was purged with N_2_ for 15 minutes; the container remained sealed at room temperature for a 20 minute cure. After gelation, the slide and adhesive mask were gently removed, excess gel was trimmed with a KimWipe, and a flow chamber (Ibidi, 80168) was affixed directly onto the coverslip to enclose the hydrogel-embedded cells for downstream imaging and filled with 2X SSC.

To reduce potential batch effects, D100 and D10000 samples were multiplexed and processed together. Samples from these experiments underwent identical primary and secondary hybridization reactions (with different barcodes) and were imaged under the same conditions, as described below.

### Par-seqFISH automated imaging

Imaging was performed using a combined widefield microscopy and automated fluidics delivery system. A widefield inverted ECLIPSE Ti2-E microscope (Nikon) equipped with a motorized stage and the NIS-Elements software suite was used for image acquisition. Focus was maintained throughout imaging using Nikon’s Perfect Focus System (PFS), and images were captured with a Prime BSI Express cMOS camera (Teledyne). All imaging was performed using a CFI Plan Apochromat 100× oil immersion objective (1.45 numerical aperture; Nikon; MRD01905). Fluorophore excitation was provided by a SOLA III FISH Light Engine (Lumencor), and fluorescence excitation/emission was filtered using standard filter cubes for DAPI, FITC (Alexa Fluor 488), TRITC (Atto 550), and Cy5 (Atto 647). Automated imaging routines were constructed using the NIS-Elements JOBS module. Fluidic flow was driven by a CETONI Nemesys S module pump equipped with a 5 mL glass syringe. Reagent delivery was managed via a Qmix valve module and the rotAXYS 360° positioning system (CETONI). The entire setup was operated using the CETONI Elements software, which was used together with the NIS-Elements JOBS to synchronize fluidics and imaging tasks, respectively.

Chambered coverslips containing gel-immobilized samples were mounted on the microscope and connected to the CETONI fluidic system. Regions of interest were selected using phase-contrast imaging. A series of sequential secondary hybridizations, washes, and imaging steps was performed to acquire barcoding and mRNA expression signals. Each secondary hybridization round included three unique 15 nt readout probes, each conjugated to either Alexa Fluor 488 (A488), Atto 550 (A550), or Atto 647 (A647). Each serial probe mixture was prepared in EC buffer composed of 10% ethylene carbonate (Sigma-Aldrich, E26258), 10% dextran sulfate (Sigma-Aldrich, D4911), 4X SSC, and 50 nM of each readout probe. Hybridization reactions were flowed into the imaging chamber and incubated with the sample for 25 minutes to allow secondary probe binding. Samples were then washed with 1 mL of 10% formamide wash buffer (10% formamide, 0.1% Triton X-100, and 2X SSC) to remove excess probes and limit nonspecific binding. This was followed by a rinse with 1 mL of 2X SSC and staining with 4′,6-diamidino-2-phenylindole (DAPI; Sigma-Aldrich, MBD0015), prepared as a 10 μg mL⁻¹ solution in 2X SSC. Imaging was performed in an anti-bleaching buffer composed of 10% (w/v) glucose, 1:100 diluted catalase (Sigma-Aldrich, C3155), 0.5 mg mL⁻¹ glucose oxidase (Sigma-Aldrich, G2133), and 50 mM Tris-HCl (pH 8) in 4X SSC. Z-stack images were acquired with 0.3 μm steps for all experiments. After each round of imaging, readout probes were stripped by flowing 1 mL of 55% wash buffer (55% formamide, 0.1% Triton X-100, and 2X SSC) through the chamber and incubating for 3 minutes, followed by a rinse with 2X SSC. This protocol of serial hybridizations, imaging, and probe stripping was repeated for 58 rounds to capture signals from 16S rRNA (for multiplexing), mRNA transcripts, background controls, and replicate hybridizations.

### Image analysis, demultiplexing, and gene expression measurement

Sequential images were registered using DAPI fluorescence to correct for mechanical shifts. Channel misalignments between fluorophores were corrected by aligning reference 16S rRNA signals across all fluorescence channels (A488, A550, and A647). Cell segmentation was performed using phase-contrast images with Cellpose 2.0^51^. A quality control step was applied to remove segmentation artifacts and autofluorescent components. Readout reaction specificity was assessed, resulting in the exclusion of 6 reactions due to insufficient specificity or aberrant signal patterns, leaving 144 high-quality genes for all downstream analyses.

For sample demultiplexing, fluorescence intensities from the background (no readouts) and 16S rRNA signals were measured within cell boundaries for each relevant readout probe to compute a signal-to-background ratio. To account for variability in baseline fluorescence across physiological conditions, a reference 16S rRNA probe was imaged in all three fluorescence channels (A488, A550, and A647). These reference and barcode signals were jointly used to assign each cell to its condition of origin. The false classification rate was estimated by quantifying the number of cells assigned to barcode combinations not used in the experiment.

To quantify gene expression, mRNA-FISH spots were identified as regional intensity maxima. Spots were filtered by an intensity threshold defined using negative control genes (P1 phage), limiting false-positive primary hybridization to <0.07%, and the no-library internal control, limiting false-positive secondary reactions to a maximum of 0.9%. An average of 0.08% of falsely detected spots is estimated across all reactions. The integrated intensity of each spot was calculated by summing pixel values within a 3×3 window centered on the local maximum. Detected spots were assigned to individual cells based on segmentation masks, and the total fluorescence signal was summed per gene per cell. Total mRNA-FISH fluorescence intensity was normalized by the number of probes designed for each gene (Table S1).

### Expression frequency analysis

We quantified the expression frequency of each gene within each density as the proportion of cells exhibiting non-zero expression. However, this measure is systematically influenced by reduced transcriptional activity at higher densities^20^. To account for these physiological dynamics, we calculated a normalized expression frequency by comparing each gene’s raw frequency to the maximum observed frequency within that specific density—representing the expected upper bound under global transcriptional constraints. For example, at an OD₆₀₀ of 2.7, the highest observed expression frequency was 72%, which was scaled to a normalized value of 100%. Thus, in this condition, a gene detected in 50% of cells was normalized to 69% of the density-specific maximum.

### Subpopulation differential expression

We analyzed differential gene expression by stratifying populations based on the expression of a target gene of interest. For each sample, cells in the top 5% and bottom 5% of expression for the target gene were designated as “gene++” and “gene−”, respectively. Gene expression values were normalized to cell size to control for total intensity differences associated with cell cycle stage. For each gene, we computed the fold change in expression between the high and low subpopulations and assessed statistical significance using a two-sided Mann–Whitney U test. *P*-values were corrected for multiple comparisons using the Bonferroni method. Genes with low expression (maximum mean fluorescence <10 units across the two subpopulations) were excluded from the analysis. Genes were considered notably differentially expressed if they exhibited a Bonferroni-adjusted *p*-value ≤ 0.01 and a fold change >2 (Table S3).

For co-expression analysis, we investigated the contribution of the top 5% of *cbpD*-expressing cells (*cbpD*++) to the expression of other individual exoproduct genes and to overall public goods production. At each population density, we identified *cbpD*++ cells and quantified the proportion of total expression for each exoproduct gene that originated from this subpopulation. To assess the collective contribution to public goods output, we also computed a metric summing the expression of all exoproduct genes within each cell, focusing on those with expression dynamics (expressed under similar densities) similar to *cbpD* (e.g., *lasA, lasB, rhlB, phzE1*). This approach enabled us to evaluate the extent to which *cbpD*++ cells drive public goods production across the population.

### Variability analysis

We found that the CV often overestimated variability in our data, primarily due to its sensitivity to outliers and an overemphasis on genes targeted by fewer probes. To mitigate this issue, we developed the TER, a complementary metric that quantifies expression variability by examining the width of the distribution among cells that express the gene. Specifically, TER is calculated as the ratio between the mean expression of the top and bottom 5% of expressing cells, thus reducing the influence of outliers. However, while TER is more robust to extreme values, it tends to underestimate variability in genes with low probe coverage—opposite to the CV. To leverage the strengths of both approaches, we combined these complementary metrics and removed several genes covered by very few probes (*rsaL, acp1, qslA, pscF*). Additionally, data-points with an average expression below 32 fluorescent arbitrary units were discarded.

CV and TER values were computed for each gene at each density in either the D100 or D10000 experiments. For TER, only cells with detectable expression of the gene were considered; these cells were ranked by expression level, and the means of the top and bottom 5% were compared. To account for potential biases due to fluorophore brightness, the merged dataset (n = 1,560 datapoints combined in D100 and D10000 experiments) was stratified by fluorophore (A488, A500, and A647). Within each fluorophore group, we modeled the relationship between log-transformed mean expression and log-transformed CV or TER using polynomial regression: degree 2 for CV and degree 3 for TER. Parameters were estimated using ordinary least squares via the *statsmodels* Python package. A 95th percentile confidence interval was estimated, and residuals (CVR and TERR) were calculated as the difference between each observed value and its predicted interval-bound value for the corresponding metric. These residuals were then standardized (z-scored) and averaged to yield a unified variability score (EVI) that balances the complementary biases of CV and TER. An EVI score of 1.82 marked the top 5% (95th percentile). Datapoints above this threshold were classified as hypervariable.

## Acknowledgements

We thank Dianne Newman, Avigdor Eldar, David Zeevi, Roy Jacobson, Irit Sherman, Nadav Hen and Zohar Persky for critically reading and commenting on the manuscript. We thank Zohar Persky for assistance and support with par-seqFISH. We also thank Avigdor Eldar for the *P. aeruginosa* PAO1 PG strain and Viviana Job for the CHA isolate. D.D. was supported by the ERC-StG programme (grant no. 101117863) and the Israel Science Foundation (personal grant 2227/22).

## Data and materials availability

All data and scripts from this study will be deposited. All other data required for reproducing the figures and conclusions of this manuscript are presented in the main text or the supplementary materials.

