## Supplemental figures 1-6 for "Single-Cell Phenotypic Heterogeneity Shapes Quorum Signaling Dynamics in *Pseudomonas aeruginosa*"

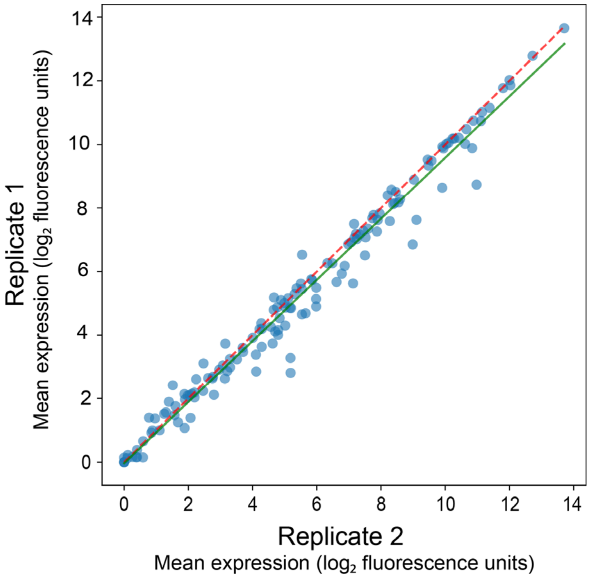


**Figure S1. Correlation between average gene expressions across biological replicates.** Log2 transformed average gene expression (total fluorescence intensity in arbitrary units) across all genes at an $OD_{600}$ of 0.24 in biological replicates (D100 vs. DB100). The dashed red line represents the identity line (X = Y). The line of best fit is shown in green (Pearson r= 0.99; $p < 10^{-10}$).

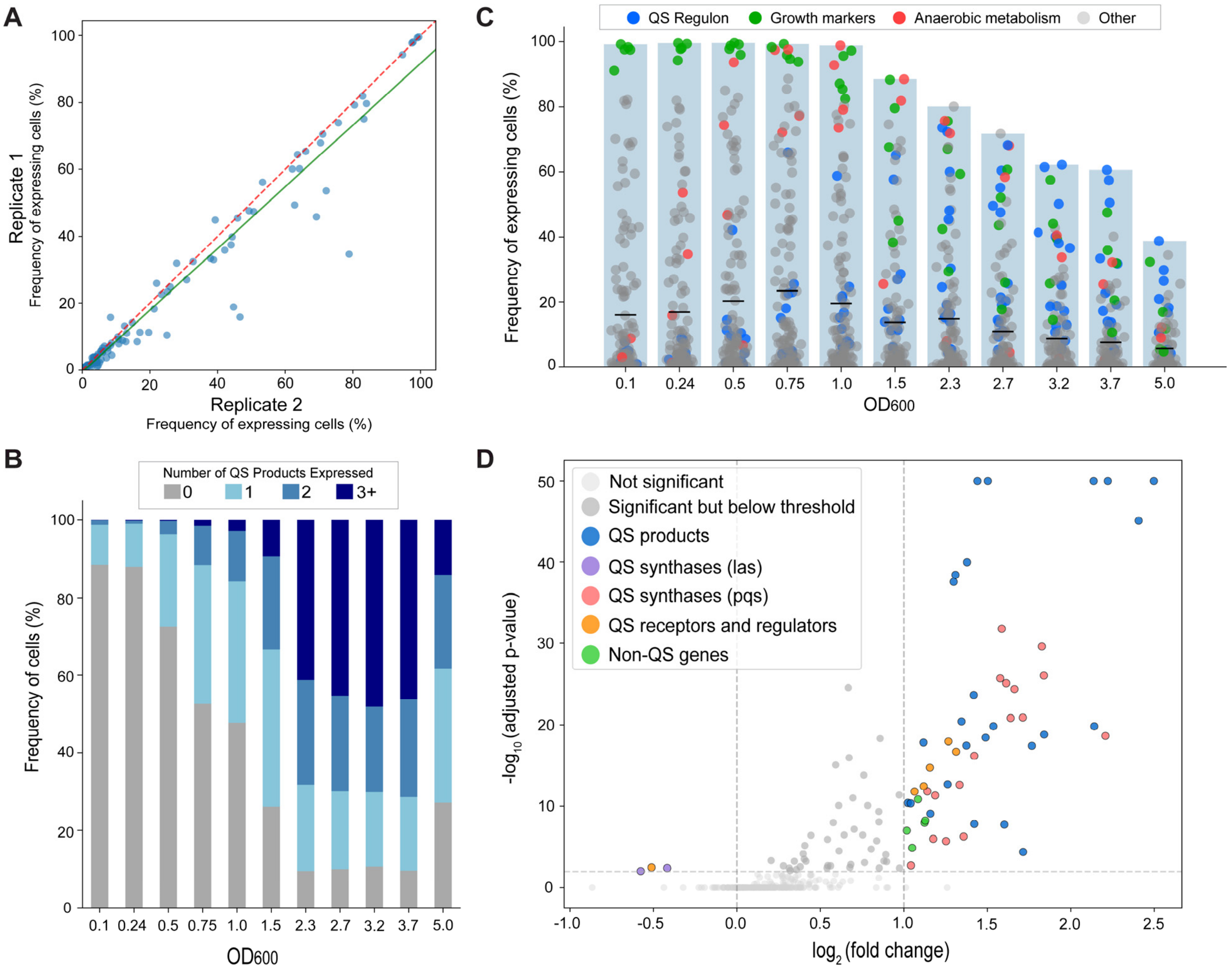


**Figure S2. Evaluating the contributions of individual bacteria to the QS response. (A)** Correlation in percentage of cells expressing each gene across biological replicates at an $OD_{600}$ of 0.24. The dashed red line represents the identity line (X = Y). The solid green line is the line of best fit (Pearson r = 0.98, $p < 10^{-10}$). **(B)** Percentage of cells expressing 0 (grey), 1 (light blue), 2 (blue), and 3 or more (dark blue) directly QS-regulated exoproducts across all densities ($OD_{600}$). **(C)** Expression frequency distributions of all measured genes across density. Bars and horizontal lines illustrate the maximum and average observed frequency per condition, respectively. Replicative capacity markers (*rpsC*, *sucC*, etc.) are marked in green, QS-regulated genes in blue, and anaerobic stress markers in red. **(D)** Volcano plot showing differential expression between subpopulations expressing high or low *cbpD* levels (top vs. bottom 5% of cells under each condition). The Y and X axes denote the Bonferroni-adjusted p-value and $log_2$ fold change, respectively. Genes differentially expressed by at least 2-fold are colored and labeled as shown in the legend.

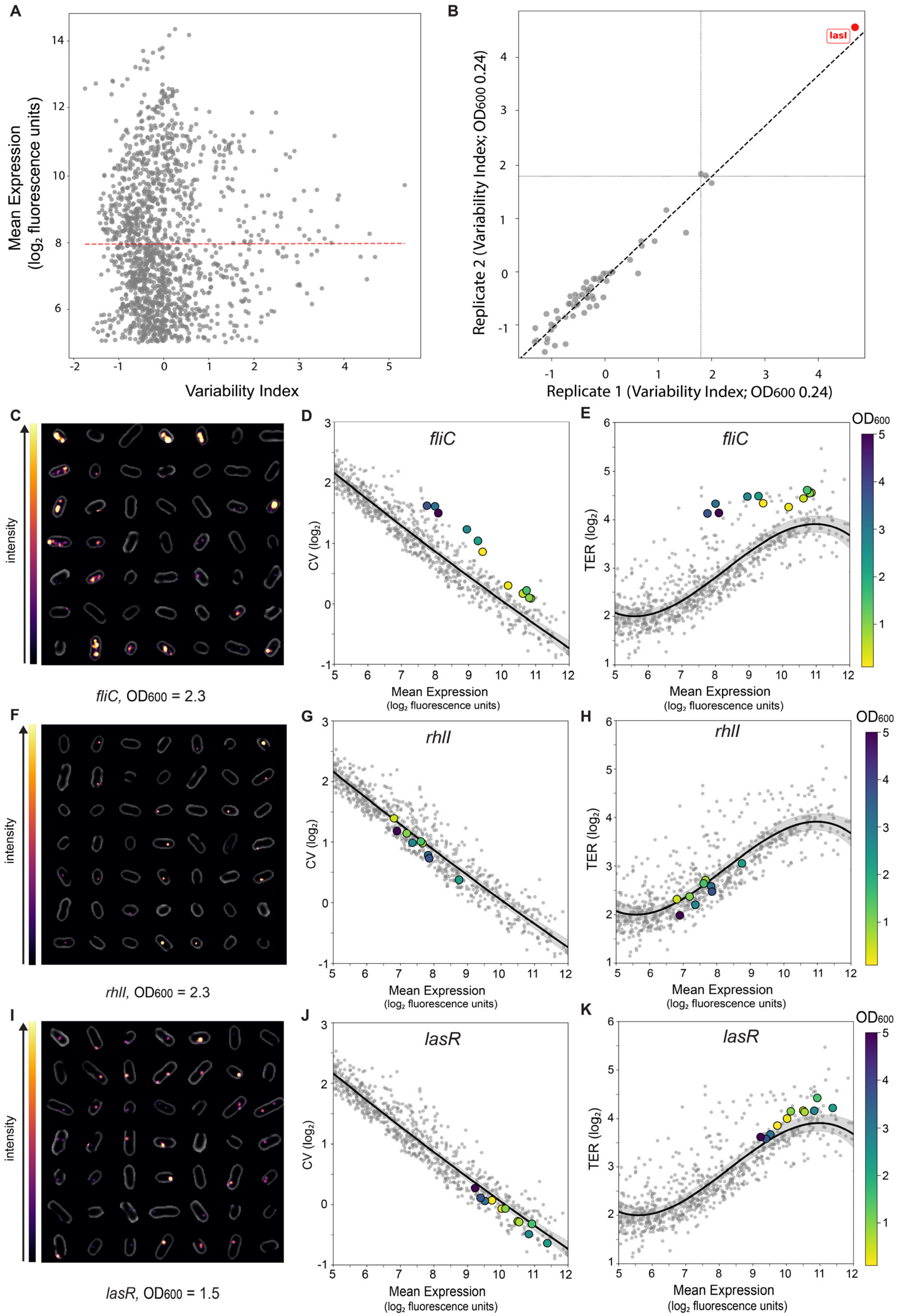

A
Mean Expression
(log2 fluorescence units)
Variability Index
B
lasI
Replicate 2 (Variability Index; OD600 0.24)
Replicate 1 (Variability Index; OD600 0.24)
C
intensity
fliC, OD600 = 2.3
D
fliC
CV (log2)
Mean Expression
(log2 fluorescence units)
E
fliC
TER (log2)
OD600
F
intensity
rhlI, OD600 = 2.3
G
rhlI
H
rhlI
I
intensity
lasR, OD600 = 1.5
J
lasR
K
lasR

**Figure S3. Expression variability index (EVI): mean-independence, replicate consistency, and gene-specific expression patterns. (A)** Correlation between $\log_2$ mean expression (arbitrary fluorescence units) and EVI across all gene-condition combinations. The dashed red line indicates the line of best fit (Pearson's r = 0.001, *p* = 0.96), demonstrating effective decoupling of mean expression and expression variability in our approach. **(B)** Correlation in EVI between biological replicates (D100 vs. DB100) at an $OD_{600}$ of 0.24, with the dashed black line indicating the line of best fit (Pearson's r = 0.98, $p < 10^{-10}$). Dotted vertical lines mark the variability threshold (EVI = 1.82). **(C)** Phase contrast images of randomly selected cells from an $OD_{600}$ of 2.3, overlaid with smFISH data for *fliC*, with signal intensity visualized using the inferno look-up table. **(E–F)** $\text{Log}_2$-transformed coefficient of variation (CV) and total expression range (TER) values plotted against $\log_2$ mean expression for all genes across all densities. Gray points represent all gene-expression measurements, while black lines show regression fits with 95% confidence intervals (gray shading). *fliC* data points are highlighted and colored by cell density as indicated. **(F–G)** Same analysis as in (E–F), highlighting *rhlI*. **(I–K)** Same analysis as in (E–F), highlighting *lasR*, with representative cell images shown in panel (I) from an $OD_{600}$ of 1.5.

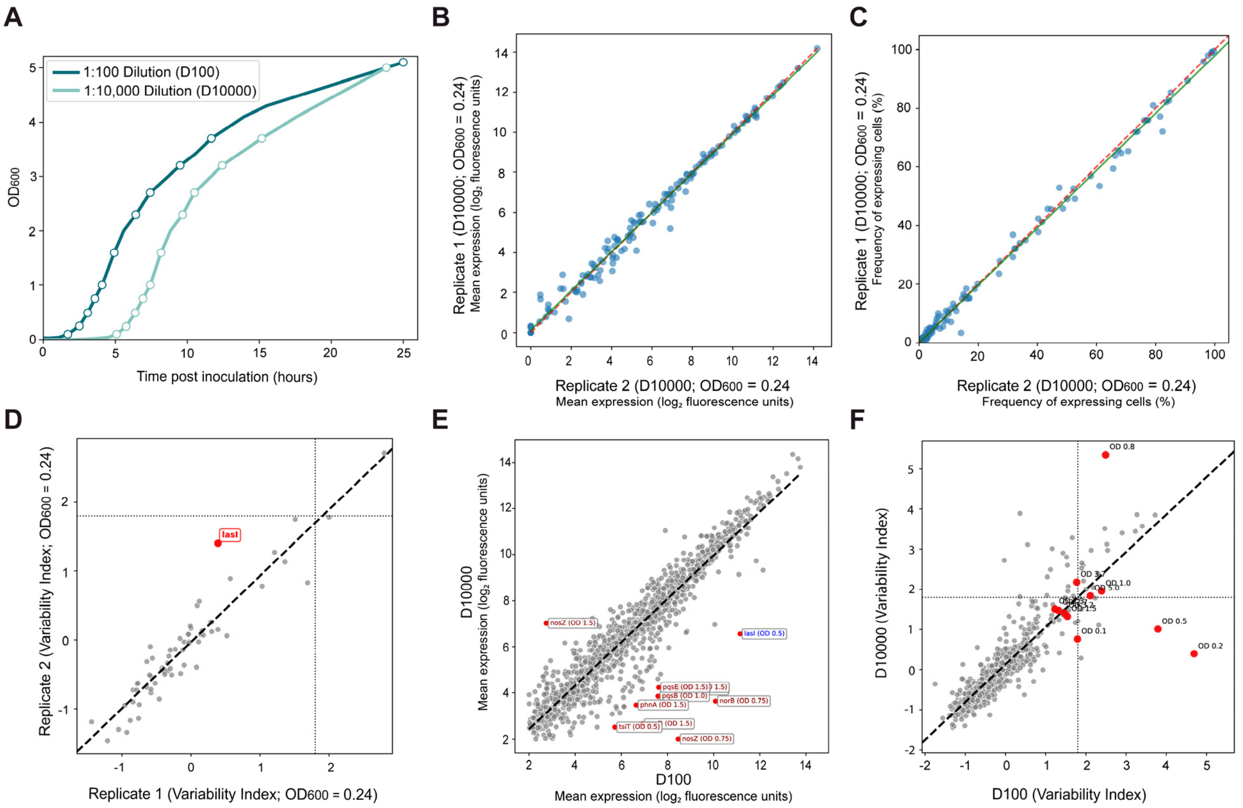


**Figure S4. Evaluating the effects of inoculum dilution on the QS response. (A)** OD$_{600}$ versus time post-inoculation (hours) for 1:100 (D100; dark turquoise) and 1:10,000 (D10000; light turquoise) dilution cultures. Sampled timepoints are indicated by circled points on the growth curves. **(B)** Log$_2$-transformed average gene expression (total fluorescence intensity in arbitrary units) across all genes at an OD$_{600}$ of 0.24 in biological replicates (D10000 vs. DB10000). The dashed red line represents the identity line ($x = y$), and the solid green line indicates the line of best fit (Pearson's $r = 0.99$; $p < 10^{-10}$). **(C)** Correlation in the percentage of cells expressing each gene across D10000 biological replicates at an OD$_{600}$ of 0.24 (Pearson's $r = 0.998$, $p < 10^{-10}$). **(D)** Correlation in EVI across biological replicates (D10000 vs. DB10000) at an OD$_{600}$ of 0.24. The dashed black line shows the line of best fit (Pearson's $r = 0.94$, $p < 10^{-10}$). Dotted vertical lines mark the EVI = 1.82 variability threshold. (**E)** Log$_2$-transformed average gene expression (total fluorescence intensity in arbitrary units) across all gene–OD$_{600}$ combinations in D100 vs. D10000. The dashed black line shows the line of best fit (Pearson's $r = 0.94$, $p < 10^{-10}$). The top 1% of differentially expressed gene–OD$_{600}$ combinations are annotated and highlighted in red. **(F)** Correlation in EVI across all gene–OD$_{600}$ combinations (gray points) in D100 vs. D10000. The dashed black line indicates the line of best fit (Pearson's $r = 0.86$, $p < 10^{-10}$). Dotted vertical lines denote the EVI = 1.82 variability threshold. *lasI* expression at various OD$_{600}$ values is marked by red circles.

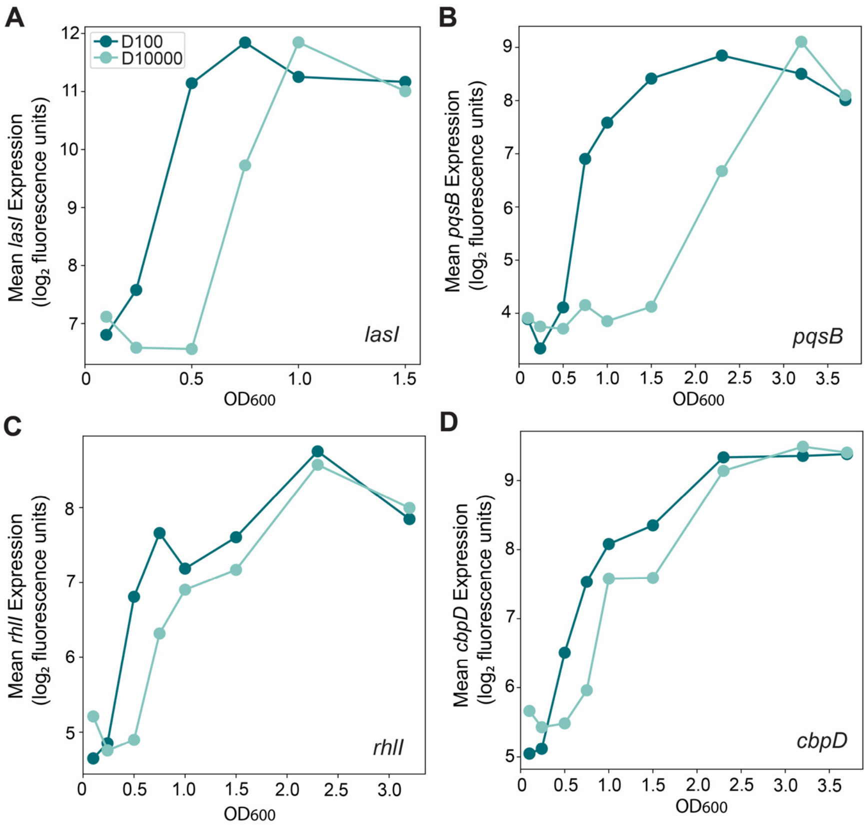


**Fig S5. Memory affects QS response dynamics. (A-D)** Log2 transformed average gene expression (total fluorescence intensity in arbitrary units) across density in D100 (dark turquoise) and D10000 (light turquoise) samples for *lasI* (A), *pqsB* (B), *rhlI* (C), and *cbpD* (D).

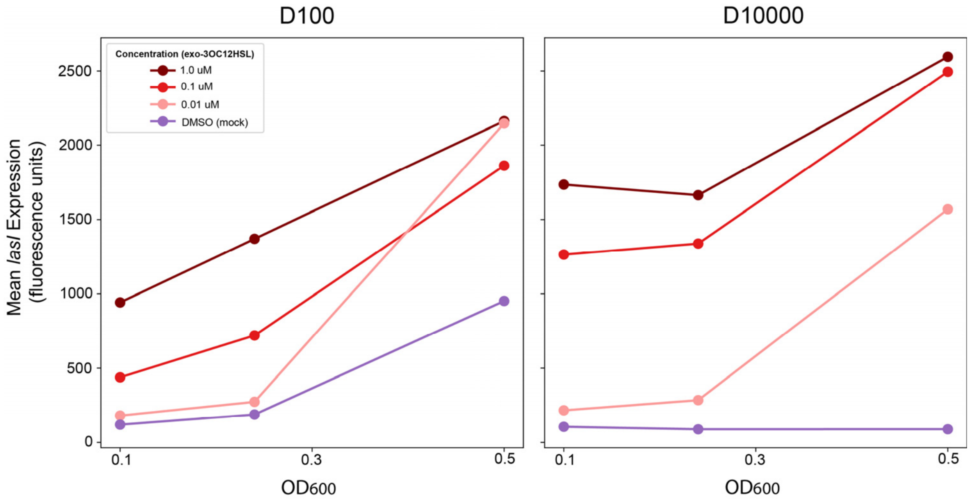


**Fig S6.** Average *lasI* expression (total fluorescence intensity in arbitrary units) across early ODs (0.1, 0.24, 0.5) during growth in either exogenous 3OC12-HSL or DMSO media for D100 and D10000 dilutions.
